# Signal-Level Witnessing of SU(1,1) Pair Dynamics in Brain Proton Spin Ensembles

**DOI:** 10.1101/2025.11.10.687594

**Authors:** Christian Kerskens

## Abstract

Non-compact symmetries such as SU(1,1) govern amplification and squeezing, yet have not been directly identified in macroscopic spin ensembles. Here we apply a symmetry-based analytical framework to previously published magnetic-resonance data acquired from proton spin ensembles in the living human brain. By reanalyzing the detected signal within this framework, we identify a non-compact SU(1,1) pair sector of the full SU(4) spin algebra whose generators carry double-quantum coherence order, and exclude compact SU(2) exchange pathways as an explanation of the observed dynamics. Although the SU(1,1) pair coherence resides in the double-quantum sector, the 45°–gradient–45° readout block converts it into a detectable signal through a specific coherence-transfer pathway.

We argue that the primary significance of the detected signal is as a witness of entry into a deep metric regime in which purely single-mode compression is no longer sufficient and cross-mode squeezing-like structure becomes necessary. In this sense, the signal functions first as a metric-regime witness and, more specifically, as a witness of non-compact pair-sector multiple-quantum coherence and squeezing. We then show that any strictly bipartite evaluation is obstructed in the high-temperature bulk-NMR setting, and that the appropriate entanglement framework is instead a macroscopic multiple-quantum-coherence witness. The observed signal features are consistent with metric-driven SU(1,1) pair dynamics, while definitive certification of many-body entanglement remains conditional on pathway-corrected calibration of the transfer coefficient and on evaluation of the corresponding separable MQC bound.

## Introduction

Recent magnetic-resonance experiments on the living human brain have revealed signals [7] whose microscopic origin remains unresolved. Within conventional nuclear-spin theory, the observed behavior is difficult to reconcile with compact SU(2) exchange dynamics, which govern bounded oscillatory evolution in dipolar-coupled ensembles. This raises the possibility that the measured signal originates instead from a non-compact dynamical sector, one that supports hyperbolic trajectories characteristic of SU(1,1) symmetry.

The central question of the present paper is not, in the first instance, whether the measured spin system is already demonstrably entangled. The more basic question is whether the detected signal indicates entry into a dynamical regime in which purely single-mode compression is no longer sufficient and cross-mode squeezing-like structure becomes necessary. In a complementary covariance-based description [6], this transition occurs when the relevant metric regime reaches the Williamson/Casimir floor and further covariance reduction must overflow into cross-mode structure. In the present spin-language description, the onset of an SU(1,1)-type pair signal is interpreted as the observable signature of that transition.

Non-compact symmetries such as SU(1,1) play a central role in quantum optics and field theory, where they govern squeezing, amplification, and pair-creation processes [9, 4, 10]. In contrast, direct observation of SU(1,1) dynamics in macroscopic spin ensembles has remained elusive. Here we reanalyze previously published data from human brain tissue [7] using a symmetry-based framework that identifies the active manifold as a non-compact SU(1,1) pair sector of the full SU(4) spin algebra.

A key subtlety is that the SU(1,1) pair operators

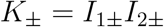

carry double-quantum (DQ) coherence order (|Δ*m*| = 2), not zero-quantum order. Nevertheless, the 45°−gradient−45° readout block employed in the experiment converts DQ pair coherence into a detectable signal through a specific coherence-transfer pathway:

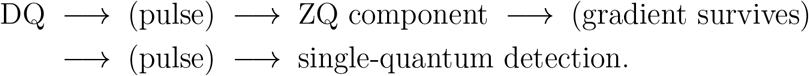

The detected signal should therefore not be interpreted as a direct observation of the algebraic sector in which the SU(1,1) generators live, but as a filtered output of that sector.

This distinction motivates the hierarchy of interpretation used throughout the paper. At the weakest level, the signal functions as a witness of entry into a non-compact metric regime, insofar as it indicates that compact single-mode exchange is no longer an adequate description. At a stronger level, the same signal functions as a witness of SU(1,1)-type pair multiple-quantum coherence and squeezing in the spin sector. Only under additional quantitative assumptions does it become a formal many-body entanglement witness.

We apply this framework to the reported brain data [7], which display a signal whose features are inconsistent with the standard signatures of compact SU(2) exchange. Our first goal is therefore to identify whether the detected signal is more naturally described as the readout-converted signature of a non-compact SU(1,1) pair sector. Our second goal is to determine the stronger condition under which the same signal may be promoted to a formal entanglement witness in the appropriate macroscopic MQC framework, and to explain why a strictly reduced two-spin evaluation fails in the high-temperature bulk setting.

The structure of the paper is as follows. Section 1 classifies the relevant subalgebras of the two-spin system and identifies the non-compact SU(1,1) pair algebra in the double-quantum sector. Section 2 derives the coherence-transfer pathway through which DQ pair coherence produces a detectable signal via the 45°–G–45° readout block. Section 3 presents the experimental features that collectively discriminate between compact SU(2) and non-compact SU(1,1) interpretations. Section 4 describes the role of the repetitive preparation train as a stroboscopic selector of the pair-correlated manifold. Section 5 then distinguishes three levels of interpretation: first, when the detected signal should be interpreted as a witness of entry into the deep metric regime; second, when it is meaningful as an MQC-based pair-sector witness; and third, under what additional conditions the same signal becomes a many-body entanglement witness via the MQC framework, resolving the pseudopure-state obstruction that defeats any strictly bipartite evaluation at room temperature. Section 6 distinguishes collective from two-subsystem realizations of the SU(1,1) generators. Section 7 discusses the results and their limitations.

## 1 Algebraic Classification of Two-Spin Coherences

We begin by classifying the dynamically closed subalgebras of the two-spin-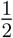 system according to coherence order. The full operator algebra is 𝔰 𝔲 (4), which decomposes into sectors of definite coherence order *p* defined by

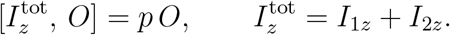

The *zero-quantum* (ZQ) sector (*p* = 0) is spanned by

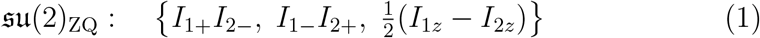

Within this space, the compact algebra

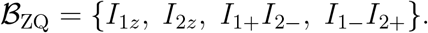

closes under commutation and generates bounded oscillatory exchange dynamics at the difference frequency Ω_−_ = *ω*_1_ − *ω*_2_ [2].

The *double-quantum* (DQ) sector (|*p*| = 2) contains the pair operators {*I*_1+_*I*_2+_, *I*_1−_*I*_2−_}. Together with the diagonal generator 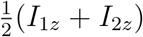 which has *p* = 0 but commutes into the DQ sector, these close to form the non-compact algebra

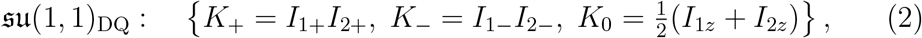

satisfying

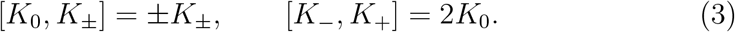

The generators *K*_±_ carry coherence order *p* =± 2 and connect the aligned states |↑↑⟩ and |↓↓⟩through pair creation and annihilation.

The crucial distinction between the two algebras lies in the geometry of their dynamics. Compact 𝔰 𝔲 (2) possesses a positive-definite Killing form and generates purely imaginary adjoint eigenvalues, leading to trigonometric time evolution (bounded oscillations). The non-compact algebra *𝔰 𝔲* (1, 1) has indefinite (Lorentzian) Killing-form signature, admits real adjoint eigenvalues, and supports hyperbolic sinh */* cosh trajectories [9, 4, 10]:

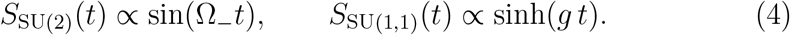

The observation that the SU(1,1) generators live in the DQ sector raises an immediate question: how can DQ pair coherence produce a detectable signal through a gradient-filtered readout that suppresses |*p* | ≠ 0 components? The answer lies in the coherence transfer pathway provided by the 45°–G–45° readout block, which we derive in the next section.

## 2 Detection Pathway: DQ Pair Coherence to Observable Signal

The 45°–G–45° readout block converts double-quantum (DQ) pair coherence into detectable single-quantum (SQ) magnetization through a coherence-transfer pathway. The purpose of this section is to establish the existence and structure of that pathway, and to define the corresponding transfer coefficient. The exact numerical calibration of the transfer coefficient is left to a full propagator simulation; the product-operator analysis given here is intended to identify the relevant transfer channels and their leading scaling.

### 2.1 Setup

Consider a two-spin state with pair coherence

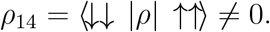

In operator language, this corresponds to the DQ pair component

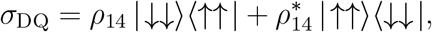

which carries coherence order *p* =± 2. The question is whether the readout block can convert this DQ component into a signal that survives the gradient and is subsequently detected as SQ magnetization.

### 2.2 Stage 1: First *R*_*x*_(*π/*4) pulse

A non-selective RF pulse with flip angle *θ* = *π/*4 about the *x*-axis acts on the one-spin basis as

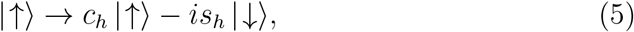

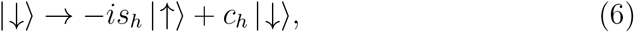

where

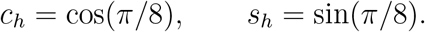

Hence

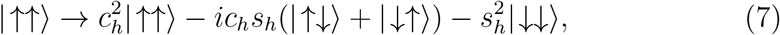

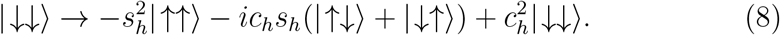

The first pulse therefore mixes the initial DQ pair coherence into components of several coherence orders. In particular, it generates a *p* = 0 contribution in the zero-quantum block,

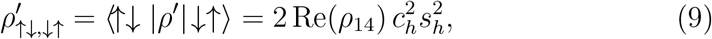

as well as population shifts in the aligned states,

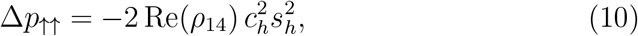

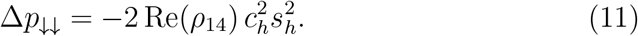

The key point is not the detailed coefficient by itself, but the fact that the first pulse transfers part of the DQ pair coherence into the *p* = 0 sector.

### 2.3 Stage 2: Gradient filter

The crusher gradient dephases all components with *p* ≠ 0. The directly DQ part of the initial pair coherence is therefore removed. What survives are precisely the *p* = 0 terms generated by the first pulse: the ZQ component Eq. (9) and the population shifts Eqs. (10) and (11).

Thus the gradient does not preserve the SU(1,1) pair coherence itself. Rather, the first pulse maps part of that coherence into a gradient-surviving intermediate sector, which is then available for conversion into detectable SQ signal by the second pulse.

### 2.4 Stage 3: Second *R*_*x*_(*π/*4) pulse and SQ detection

The second 45° pulse acts on the surviving *p* = 0 terms and converts them into SQ magnetization. Two classes of contribution are relevant:

i. The ZQ term 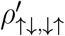 from Eq. (9). Under *R*_*x*_(*π/*4), the off-diagonal element connecting |↑↓⟩ and |↓↑⟩ is partially converted into single-quantum coherences (*p* = ±1). Specifically, the rotation mixes |↑↓⟩ and with and, generating terms proportional to *I*_1+_, *I*_2+_, etc., which contribute to the detected transverse magnetization.
ii. The population shifts Δ*p*_↑↑_ and Δ*p*_↓↓_ from Eqs. (10) and (11). Population differences along the *z*-axis are converted into transverse magnetization by the 45° tipping pulse, contributing signal proportional to sin 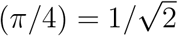 per spin.

Both classes are linear in the initial *ρ*_14_ and therefore contribute linearly to the measured transverse signal. The full measured response is the coherent sum of these conversion channels, together with any further contributions introduced by finite pulse duration, relaxation during the readout, and RF inhomogeneity.

### 2.5 Definition of the transfer coefficient

The net effect of the complete readout block is therefore summarized by

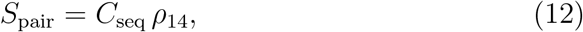

where *C*_seq_ is the effective transfer coefficient for the DQ-pair-sector pathway. This coefficient includes the full DQ → *p* = 0 →SQ conversion, including interference between the ZQ and population channels.

The product-operator calculation above establishes two facts:

a. the transfer coefficient is nonzero, so DQ pair coherence can contribute to the detected signal;
b. the transfer is suppressed relative to a direct ZQ→SQ conversion, because it proceeds through an intermediate pathway rather than through direct gradient survival of the original coherence. Using

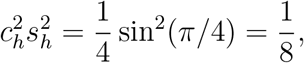

the leading pathway estimate is of order

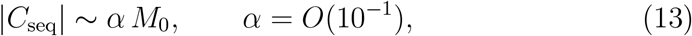

with the simplest pathway estimate giving *α* ∼ 1*/*8. This number should be regarded as an order-of-magnitude indicator, not as a final experimental calibration.

### 2.6 Interpretive consequence

The main consequence for the rest of the paper is qualitative but important: the effective transfer coefficient for the pair-sector pathway is expected to be substantially smaller than the coefficient that would apply to a directly detected zero-quantum coherence. Therefore, for a fixed measured signal amplitude, the inferred pair-sector coherence |*ρ*_14_| is correspondingly larger once the DQ →SQ conversion cost is taken into account.

A definitive numerical value of *C*_seq_ requires a full simulation of the actual pulse sequence, including finite pulse duration, relaxation during the readout, off-resonance effects, and RF inhomogeneity. The witness analysis of Section 5 therefore uses Eq. (12) formally and treats the exact calibration of *C*_seq_ as an open quantitative problem.

## 3 SU(2) vs. SU(1,1) Signatures

We now examine whether the measured signal [7] can be explained within a compact SU(2) exchange picture or instead requires a non-compact SU(1,1) pair-mode description. The argument is cumulative: no single feature is decisive in isolation, but taken together they strongly disfavor a compact interpretation. Table 1 summarizes the comparison; the individual features are discussed below.

**Table 1.** Discrimination between compact SU(2) and non-compact SU(1,1) interpretations of the measured signal. Each row lists an observable feature together with the prediction of each algebra.

| Feature | Compact SU(2) | Non-compact SU(1,1) | Observed |
| --- | --- | --- | --- |
| Preparation | Requires population imbalance $(I_{1z} - I_{2z}) \neq 0$ | Pair creation from balanced state via $K_{\pm}$ | Signal under balanced prep |
| Time dependence | Bounded: $\sin(\Omega_- t)$ | Hyperbolic: $\sinh(gt)$ | Non-oscillatory onset |
| Refocusing frequency | Difference: $\Omega_- = \omega_1 - \omega_2$ | Pair: $\Omega_+ = \omega_1 + \omega_2$ | Refocuses at $\Omega_+$ |
| Echo parity | Refocuses every echo | Revives every second echo | Even-echo structure |
| $M_0$ scaling | Quadratic (collective) or bounded | Linear (bipartite pair) | Linear |
| Magic angle | Suppressed | Suppressed | Suppressed |
| Coherence pathway | $\rho_{23}: \vert \uparrow\downarrow\rangle \leftrightarrow \vert \downarrow\uparrow\rangle$ | $\rho_{14}: \vert \uparrow\uparrow\rangle \leftrightarrow \vert \downarrow\downarrow\rangle$ | Consistent with $\rho_{14}$ |
| Reservoir coupling | Classical intensity modulation | Phase-locked pair via $[K_0, K_{\pm}] = \pm K_{\pm}$ | Coherent pair signal |

### Balanced populations

Following presaturation, the longitudinal magnetization is approximately balanced, ⟨*K*_0_⟩ ≈ 0. Compact SU(2) exchange generated by *I*_1+_*I*_2−_ + *I*_1−_*I*_2+_ requires a population imbalance proportional to (*I*_1*z*_ − *I*_2*z*_) to produce ZQ coherence. The observation of a finite signal under approximately balanced preparation therefore disfavors a compact SU(2) exchange mechanism.

### Absence of an evolution delay

The signal appears directly after the 45°–G–45° block without an additional free-evolution interval. In a compact SU(2) picture, ZQ coherence requires finite-time exchange evolution. The immediate appearance is not the expected behavior for a standard compact exchange pathway.

### Echo-parity behaviour

The signal alternates in sign and revives every second echo, consistent with a sign reversal of the SU(1,1) quadrature under the crusher sequence. Compact SU(2) echoes refocus at each echo time and do not naturally produce this parity structure.

### Frequency refocusing

The observed echo refocuses at the pair frequency Ω_+_ = *ω*_1_ + *ω*_2_, whereas compact SU(2) ZQ exchange is associated with Ω_−_ = *ω*_1_ − *ω*_2_. This favors a pair-mode interpretation.

### Amplitude scaling

The amplitude scales linearly with *M*_0_. As shown in Section B, quadratic scaling is the natural expectation for collective single-manifold SU(1,1) squeezing, whereas linear scaling is consistent with selective pair-mode coupling between distinguishable subsystems.

### Coherence-pathway identity

The detected signal traces back to the density-matrix element *ρ*_14_, which connects | ↑↑⟩ and | ↓↓⟩. through the SU(1,1) pair operators, as established by the pathway analysis of Section 2. Compact SU(2) exchange acts on the *ρ*_23_ sector connecting |↑↓⟩ and |↓↑⟩.

### Magic-angle behaviour

The signal vanishes at the magic angle *θ*_*m*_ = arccos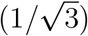confirming dependence on anisotropic dipolar coupling. While not unique to SU(1,1), this is consistent with a two-spin pair mode and, combined with the preceding features, supports the non-compact assignment.

### Reservoir coupling structure

A purely classical reservoir can modulate intensities but cannot generate a phase-locked pair amplitude ⟨*I*_1+_*I*_2+_⟩. The observed coherence is consistent with coupling through *K*_0_ via the SU(1,1) commutation relations [*K*_0_, *K*_±_] =± *K*_±_.

Taken together, these features strongly disfavor a compact SU(2) zero-quantum exchange scenario and support the interpretation that the measured signal arises from a non-compact SU(1,1) pair sector.

## 4 The Repeated Preparation Train as a Stroboscopic Selector

A central feature of the experiment is that the relevant preparation is not a single 45°–G–45° block but a repetitive train of such blocks. This repetition establishes a stroboscopic dynamical environment whose role should be stated carefully in light of the algebraic structure identified in Section 1.

The SU(1,1) pair operators *K*_±_ = *I*_1±_*I*_2±_ generate DQ coherence (coherence order |*p*| = 2). Between preparation blocks, any active pair-creation process populates {| ↑↑⟩, | ↓↓⟩}the, subspace and generates DQ pair correlations. Each 45°–G–45° block then acts in two capacities:

i. *Readout* : it converts a fraction of the current DQ pair coherence into detectable SQ magnetization through the pathway derived in Section 2;
ii. *Re-preparation*: it projects the system back into a restricted manifold by destroying coherences that do not survive the gradient filter.

The stroboscopic picture therefore describes repeated cycles of pair creation → partial readout → re-preparation.

The repetitive train does not by itself guarantee a large pair-sector signal. Its primary role is to create a reduced dynamical environment in which the DQ pair sector survives repeated filtering and in which non-compact pair dynamics are not overwhelmed by conventional single-quantum pathways. Observable signal still requires a squeezing-like gain process that transfers weight from the longitudinal sector *K*_0_ into the pair sector *K*_±_.

This viewpoint can be formalized as a discrete stroboscopic map. Let *ρ*_*n*_ denote the reduced state immediately before the *n*th block. One cycle defines an effective map

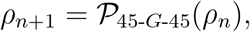

whose action on the pair-sector expectation values takes the form

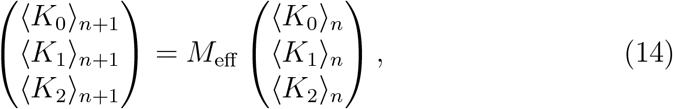

Where 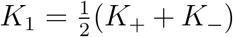 and 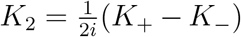 are the Hermitian quadratures, and *M*_eff_ encodes the coarse-grained action of one block together with any active pair-generating dynamics. If *M*_eff_ has boost-like non-compact character, repeated application amplifies the pair quadrature in the manner expected of SU(1,1) dynamics.

It is important to note that the gradient does not “preserve” the SU(1,1) sector: it destroys DQ coherence directly. Rather, the pulse–gradient–pulse sandwich provides an asymmetric coherence transfer that converts DQ pair correlations into a selectively detectable signal (Section 2). Between blocks, the pair-creation process regenerates DQ coherence, which is then partially read out by the next block. The cumulative effect is a stroboscopic amplification cycle whose net character is determined by *M*_eff_.

A candidate microscopic origin of the required non-compact gain is identified in a complementary covariance-based analysis [6], where cross-mode covariance overflow beyond the Williamson/Casimir floor provides a squeezing-like mechanism coupling the same spin sectors. The present paper addresses the signal-level consequences of the non-compact dynamics; the covariance-based analysis addresses the candidate origin of the gain.

## 5 From Metric-Driven Squeezing to Entanglement Witnessing

We now distinguish three levels of interpretation for the detected signal. The first is whether the signal should be interpreted as evidence that the system has entered the deep boundary regime in which purely single-mode compression is no longer sufficient and cross-mode squeezing becomes algebraically necessary. The second is whether the detected coherence should be interpreted as a witness of non-compact pair-sector squeezing. The third, and strongest, is whether the same signal becomes an entanglement witness. The first is the primary metric question, the second is the squeezing question, and the third is a stronger, subsequent condition.

In this hierarchy, the detected signal is first a witness of entry into a non-compact metric regime, then a witness of SU(1,1)-type pair squeezing, and only under stronger assumptions an entanglement witness.

### 5.1 Signal as a witness of the deep boundary regime

The complementary covariance-based analysis [6] identifies the deep boundary regime as the point at which single-mode compression reaches the Williamson/ Casimir floor and further covariance reduction must overflow into cross-mode structure. In the present spin-language description, the onset of a non-compact SU(1,1) pair-sector signal is interpreted as the observable signature of that transition. The detected signal therefore functions first as a witness that the system has entered a metric regime in which cross-mode squeezing-like structure has become necessary.

This metric interpretation is weaker than an entanglement claim. It requires only that the detected signal be inconsistent with compact single-mode exchange and consistent with non-compact pair-sector squeezing. The evidence summarized in Section 3 — refocusing at the pair frequency Ω_+_, even-echo structure, immediate onset, balanced preparation, and incompatibility with compact SU(2) exchange — supports precisely this weaker claim.

### 5.2 Signal–density-matrix relationship

The pathway analysis of Section 2 establishes the existence and direction of the DQ→detectable-signal conversion and defines the corresponding transfer coefficient *C*_seq_. At the present stage, this coefficient is known formally but not yet numerically calibrated by a full simulation of the experimental sequence. Accordingly, we write

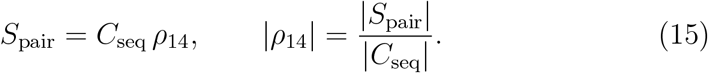

Here *S*_pair_ denotes the component of the measured signal assigned to the SU(1,1) pair pathway; experimentally, this is identified with the dominant refocusing component at Ω_+_.

The same readout block also converts any residual ZQ coherence *ρ*_23_ into SQ signal, but through a different pathway with a different transfer coefficient. The two contributions are distinguished, in principle, by their distinct refocusing frequencies:

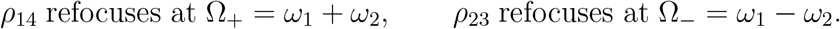

The experimental observation that the dominant signal refocuses at Ω_+_ therefore supports the working assignment

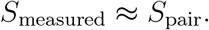

At this stage, Eq. (15) already supports a squeezing interpretation: a nonzero pathway-corrected pair-sector signal implies nonzero pair coherence and therefore signals departure from a purely compact-exchange description.

### 5.3 The bipartite thermal obstruction

If one attempts to evaluate entanglement strictly within a reduced two-spin X-state model, one encounters a severe thermodynamic obstruction. In a high-temperature bulk NMR ensemble (such as living tissue at 310 K), the density matrix is heavily dominated by the identity. The anti-aligned populations are approximately uniformly distributed,

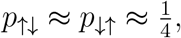

so the separable boundary for the two-qubit concurrence [11] is of order 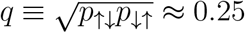.

However, macroscopic NMR signals are calibrated against the thermal equilibrium magnetization *M*_0_, which scales with the tiny thermal polarization *ϵ* = ℏ*ω/*(2*k*_*B*_*T*) 10^−5^ at clinical field strengths. Even if the pathway-corrected pair signal appears macroscopic relative to equilibrium (|*S*_pair_| *∼ M*_0_), the absolute magnitude of the off-diagonal density-matrix element is only|*ρ*_14_|∼*ϵ*∼10^−5^.

Inserting these physical scales into a bipartite concurrence formula yields

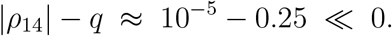

This is the well-known pseudopure-state obstruction of liquid-state NMR [1]: high-temperature macroscopic ensembles lie deep inside the separable ball, because the identity-dominated background sets *q* ≈ 1*/*4 regardless of the signal. The obstruction is not a technical limitation that can be overcome by better calibration; it is a structural feature of the reduced two-spin description applied to room-temperature bulk NMR.

Consequently, treating the signal as a strictly isolated two-spin witness creates a fundamental contradiction: the collective macroscopic signal cannot satisfy a bipartite bound that was derived for a single isolated pair embedded in a maximally mixed background. The resolution is to recognize that the detected signal is not the signature of a single isolated pair, but the collective coherence of a macroscopic ensemble — and to evaluate it within a framework designed for that setting.

### 5.4 Macroscopic many-body entanglement via MQC

To resolve the bipartite thermal obstruction, one must abandon the strictly reduced two-spin picture and treat the signal for what it physically is: a collective multiple-quantum coherence (MQC) intensity arising from a macroscopic spin ensemble.

The detected signal belongs naturally to an MQC framework. Although the SU(1,1) generators *K*_±_ = *I*_1±_*I*_2±_ reside in the DQ sector, the 45°–G–45° readout converts this pair coherence into a detected signal through the pathway derived in Section 2. The measured response is therefore a readout-converted signature of a definite collective coherence sector, rather than a single off-diagonal element of an isolated pair.

This perspective is important because MQC-sector intensities can serve directly as many-body entanglement witnesses in macroscopic, high-temperature ensembles without requiring pseudopure states or bipartite isolations. In particular, Gärttner, Hauke, and Rey showed that MQC spectra provide experimentally accessible lower bounds on the quantum Fisher information (QFI), and thereby witness genuine many-body entanglement [3]. In their framework, the generation of MQC intensity at coherence order *m* (in the present case, the DQ pair sector with|*m*| = 2) is evaluated against the maximum variance achievable by a fully separable thermal state. The witness therefore takes the form of a ratio — detected collective coherence intensity versus separable thermal bound — rather than the absolute subtraction |*ρ*_14_|™ *q* that fails in the bipartite setting.

Two points of caution are in order. First, the Gärttner–Hauke–Rey framework was developed for compact SU(2) collective generators with bounded spectra, whereas the present SU(1,1) generators are non-compact and unbounded. This adaptation is nontrivial: the Gärttner–Hauke–Rey construction carries over at the level of witness strategy, but the explicit separable bound must be rederived for the present setting. Second, treating the pathway-corrected signal amplitude as proportional to the relevant DQ-sector MQC intensity requires the same transfer-coefficient calibration discussed in Section 2; the MQC framework resolves the thermodynamic obstruction but does not eliminate the need for quantitative pathway calibration.

### 5.5 Formal macroscopic signal-level witness

In the MQC framework, a system possesses genuine many-body entanglement if the normalized DQ coherence intensity exceeds the classical variance bound of a separable state. By treating the pathway-corrected signal amplitude as proportional to the DQ-sector MQC intensity,

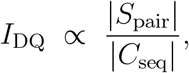

the corresponding macroscopic entanglement witness takes the form

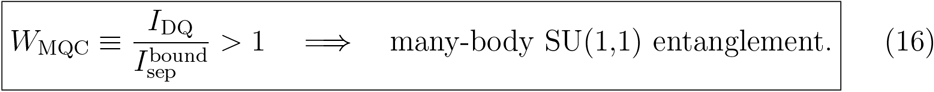

Here 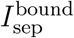 denotes the maximum DQ-sector MQC intensity achievable by a fully separable thermal ensemble under the same experimental conditions.

Equation Eq. (16) should be read as a conditional upgrade rule: once the signal has already been interpreted as a witness of metric-driven pair squeezing (Section 5.1), this stronger inequality specifies when the same signal also qualifies as an entanglement witness. The formulation as a ratio rather than an absolute subtraction is what resolves the thermodynamic obstruction: the separable bound 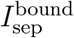 is evaluated for the same macroscopic thermal ensemble and therefore absorbs the identity-dominated background that defeats the bipartite approach.

The evaluation of 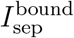 for the present non-compact SU(1,1) setting is not carried out in this paper and constitutes a concrete target for future work. What the present analysis establishes is the formal witness structure and the hierarchy of interpretations:

i. the detected signal is first a metric-regime witness (inconsistency with compact SU(2) exchange);
ii. it is second an SU(1,1)-pair-sector MQC/squeezing witness (nonzero collective DQ coherence intensity);
iii. it becomes a many-body entanglement witness only when 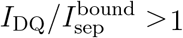 is quantitatively established.

### 5.6 Status of the numerical calibration

For the data of Ref. [7], the measured signal amplitude is of order

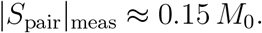

A further correction arises from transverse dephasing during the readout interval. If the detected component decays with

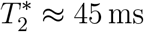

and the relevant delay before detection is of the same order,

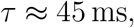

then the measured amplitude is suppressed by approximately

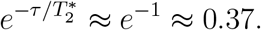

The corresponding decay-corrected pair-sector signal is therefore

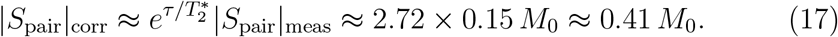

The pathway analysis of Section 2 indicates that the effective transfer coefficient should be smaller than the coefficient appropriate to a directly detected zero-quantum pathway, because the DQ pair coherence must first be routed through a gradient-surviving intermediate sector before being converted to SQ signal. A leading product-operator estimate suggests a coefficient of the form

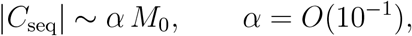

with the simplest pathway estimate giving *α* ∼1*/*8. This number should be regarded only as an order-of-magnitude indicator, not as a final sequence calibration.

If the leading-order estimate is inserted directly into Eq. (15) using the decay-corrected amplitude Eq. (17), the inferred single-pair coherence exceeds the physical bound |*ρ*_14_| ≤ 1/2 by a wide margin. As discussed in Section 5.3, this is not surprising: the signal is collective, and its interpretation requires the MQC framework rather than a single-pair density-matrix element. The excess confirms that the detected signal reflects coherent contributions from multiple effective pairs, consistent with the macroscopic MQC interpretation of Section 5.4.

A definitive quantitative evaluation of Eq. (16) therefore requires:

i. a full numerical simulation of *C*_seq_ for the actual pulse sequence, including finite-pulse effects and relaxation during the readout;
ii. computation of 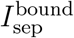 for the non-compact SU(1,1) DQ sector, adapting the Gärttner–Hauke–Rey QFI framework to generators with unbounded spectra;
iii. an empirical separation of pair-sector and compact-exchange contributions using their distinct refocusing frequencies;
iv. an explicit mapping between the detected DQ-derived signal and the corresponding MQC intensity entering Eq. (16).

These constitute the concrete calibration program required to promote the present formal witness from a metric/squeezing diagnostic to a quantitatively testable entanglement criterion.

#### Remarks

(i) The signal should be interpreted in stages: first as a metric-regime witness, then as an SU(1,1)-pair-sector MQC/squeezing witness, and only under the stronger quantitative condition 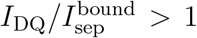 as a many-body entanglement witness. (ii) The MQC witness framework is valid in open conditions and does not require isolation of a single spin pair; it is designed precisely for the macroscopic, high-temperature setting in which the bipartite approach fails. (iii) In strictly collective realizations confined to a single manifold, the detected MQC intensity diagnoses SU(1,1) nonclassicality rather than inter-subsystem entanglement. A variance-based SU(1,1) criterion such as 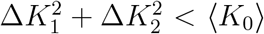 is then the more appropriate witness [8].

## 6 Collective vs. Two-Subsystem Realization

Whether the detected SU(1,1) nonclassicality constitutes entanglement between distinguishable subsystems depends on how the generators are physically realized. This question is orthogonal to the bipartite thermal obstruction discussed above: it concerns the tensor-product structure of the Hilbert space, not the thermodynamic scaling of the density matrix.

For a single-mode or collective embedding,

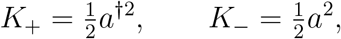

violation of an SU(1,1) witness signals internal squeezing within one collective Hilbert space. For a two-mode realization,

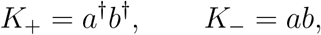

the same class of witnesses probes correlations between separate subsystems *A* and *B* [5, 8]. In the MQC framework, this distinction determines whether the witnessed entanglement is between collective modes of two distinguishable spin populations or within a single collective degree of freedom.

In the present setting, the operators *I*_1±_ and *I*_2±_ act on two physically distinct spin subspaces, so *K*_±_ = *I*_1±_*I*_2±_ generates a bipartite SU(1,1) realization. Several experimental features support this two-subsystem assignment:

a. The signal vanishes at the magic angle, identifying it as arising from anisotropic coupling between spatially distinct spin groups.
b. The amplitude scales linearly with *M*_0_ rather than quadratically, consistent with selective pair-mode coupling between distinguishable subsystems (Section B) rather than collective single-manifold squeezing.
c. The signal refocuses at the pair frequency Ω_+_ = *ω*_1_ + *ω*_2_, characteristic of a pair mode linking two distinguishable components.
d. Independent calibration of 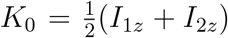 resolves additive contributions from two distinguishable components.

Under the two-subsystem assignment, violation of the MQC witness Eq. (16) would be appropriately interpreted as entanglement between the two collective spin populations rather than merely nonclassicality within a single collective mode.

## 7 Discussion

We have reanalyzed previously published magnetic-resonance data from the living human brain within a symmetry-based framework and found that the observed signal is more naturally described by a non-compact SU(1,1) pair sector than by compact SU(2) exchange. The SU(1,1) pair operators *K*_±_ = *I*_1±_*I*_2±_ generate double-quantum coherence connecting the aligned states |↑↑⟩ and |↓↓⟩. A central result of the paper is that this double-quantum pair coherence need not be directly observable in order to be experimentally relevant: the 45°–G–45° readout block provides a specific coherence-transfer pathway through which pair-sector coherence is converted into a detectable signal. The measured signal is therefore interpreted not as a direct zero-quantum excitation of the SU(1,1) generators, but as a filtered output of the double-quantum pair sector.

The primary interpretive contribution of the paper is a three-level hierarchy for the detected signal:

i. *Metric-regime witness*. The signal is inconsistent with any compact SU(2) exchange description (as established by the cumulative evidence of Section 3) and is therefore interpreted as a witness of entry into a non-compact regime in which cross-mode squeezing becomes algebraically necessary.
ii. *MQC/squeezing witness*. The detected pair-sector signal is meaningful as a witness of nonclassical many-body coherence structure within the MQC framework [3], even before one imposes a reduced two-spin model.
iii. *Many-body entanglement witness*. Under the formal condition 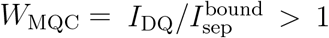, the same signal would certify many-body entanglement, bypassing the pseudopure-state obstruction that defeats any strictly bipartite evaluation at room temperature.

This hierarchy clarifies the relationship between the present paper and the complementary covariance-based analysis [6]: that work identifies when the boundary metric forces cross-mode squeezing to begin, whereas the present paper identifies how that squeezing appears in the spin signal and under what stronger conditions it can be promoted to an entanglement witness.

A key insight of the analysis is that a strictly bipartite reduced two-spin witness is thermodynamically impossible in room-temperature bulk NMR (Section 5.3). The identity-dominated background sets the separable bound at *q*≈ 1*/*4, while the absolute magnitude of any single-pair coherence is of order *ϵ*∼ 10^−5^. This is the well-known pseudopure-state obstruction [1], and it cannot be resolved by better calibration within a bipartite framework. The resolution is to recognize that the detected signal is a collective MQC intensity and to evaluate it within a framework designed for macroscopic ensembles [3]. The MQC witness formulation as a ratio 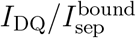 absorbs the identity-dominated background into the separable bound itself, making the comparison physically meaningful.

The pathway perspective helps organize the experimental evidence. The combination of balanced preparation, immediate signal onset, refocusing at the pair frequency Ω_+_ = *ω*_1_+*ω*_2_, even-echo structure, and magic-angle dependence is difficult to reconcile with a conventional compact SU(2) exchange interpretation. Taken together, these features favor a non-compact pair-mode description in which the detected signal inherits its structure from the SU(1,1) sector through the readout transfer pathway.

A further conceptual point concerns the role of the repetitive 45°–G–45° preparation train. The train does not itself preserve double-quantum pair coherence through the gradient; rather, it acts stroboscopically, repeatedly reading out a fraction of the pair sector and re-preparing the system for the next cycle. On this view, the observed signal reflects repeated cycles of pair generation, partial conversion into detectable coherence, and re-preparation. The present paper should therefore be read alongside the complementary covariance-based analysis [6]: that work identifies when the boundary metric forces cross-mode squeezing to begin, whereas the present paper identifies how that squeezing appears in the spin signal.

The present paper does not yet supply a definitive numerical certification of witness violation at level (iii). The leading-order pathway estimate of the transfer coefficient establishes the existence and direction of the DQ→ZQ→SQ conversion, but a full simulation is needed for quantitative calibration. More fundamentally, the separable bound 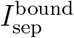 for the non-compact SU(1,1) DQ sector has not yet been computed; adapting the Gärttner–Hauke–Rey QFI framework to generators with unbounded spectra is a concrete theoretical target. The witness should therefore be regarded, at the present stage, as formally derived and experimentally targeted, but not yet numerically closed.

This status is nonetheless scientifically useful. The *T* ^∗^-corrected signal amplitude (≈ 0.41 *M*_0_) is a substantial fraction of the equilibrium magnetization, indicating that the underlying pair-sector coherence is macroscopic rather than a small perturbation. The pathway analysis implies that the effective transfer coefficient is smaller than it would be for a direct zero-quantum calibration, which further increases the inferred collective pair coherence. These features are consistent with a macroscopic SU(1,1) squeezing process and motivate the quantitative evaluation of the MQC witness.

The interpretation of the witness also depends on the physical realization of the SU(1,1) generators (Section 6). In a collective single-manifold realization, violation would diagnose SU(1,1) nonclassicality rather than inter-subsystem entanglement. The experimental features — approximately linear scaling with *M*_0_, pair-frequency refocusing, and anisotropic coupling signatures — support a two-subsystem interpretation, under which the MQC witness diagnoses entanglement between two distinguishable collective spin populations.

Within these limits, the present paper supports three claims. First, the detected signal is more consistently described as the readout-converted signature of a non-compact SU(1,1) pair sector than as a compact SU(2) exchange signal. Second, the same signal is already meaningful as a metric-regime and MQC/squeezing witness for nonclassical collective structure. Third, the macroscopic MQC framework provides a formal many-body entanglement witness whose numerical evaluation now depends on a clearly defined calibration program rather than on an unresolved conceptual gap.

Future work should focus on three priorities: (i) a full numerical simulation of the 45°–G–45° propagator, including finite-pulse effects, relaxation, and interference between transfer pathways, in order to determine *C*_seq_ quantitatively; (ii) computation of 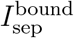 for the non-compact SU(1,1) DQ sector, adapting the QFI-based MQC witness framework to generators with unbounded spectra; and (iii) sequence-variation experiments that separate pair-sector and compact-exchange contributions through their distinct frequency and echo signatures. These steps would allow the formal witness derived here to be turned into a quantitatively testable many-body entanglement criterion for the SU(1,1) pair sector.

### A Hyperbolic evolution and compact algebras

We show that a ZQ-detected observable with hyperbolic time dependence cannot arise from compact SU(2) exchange alone. Under compact 𝔰 𝔲 (2)_ZQ_ evolution generated by *H* = *J*(*I*_1+_*I*_2−_ +*I*_1−_*I*_2+_), any expectation value ⟨*O*(*t*)⟩ with *O* ∈ 𝔰 𝔲 (2)_ZQ_ satisfies

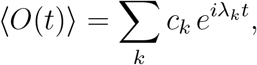

where *λ*_*k*_ are eigenvalues of the adjoint action of *H* on 𝔰 𝔲 (2). Since 𝔰 𝔲 (2) is compact, the Killing form is negative definite, and all adjoint eigenvalues are purely imaginary. The time evolution is therefore a finite sum of oscillatory terms; sinh(*gt*) growth is impossible within the compact algebra.

For the non-compact algebra *𝔰* 𝔲 (1, 1), the Killing form has Lorentzian signature (+,™, ™), and the adjoint representation admits real eigenvalues ± *g*. The corresponding time evolution includes cosh(*gt*) and sinh(*gt*) terms, producing the hyperbolic growth characteristic of squeezing and parametric amplification.

### B Scaling of SU(1,1) signal with *M*_0_

For a collective single-manifold SU(1,1) realization with 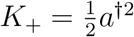 the pair amplitude scales as 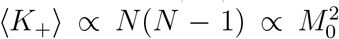, where *N* is the total spin number and *M*_0_∝ *N*. This quadratic scaling is characteristic of collective squeezing within a single mode.

For a bipartite two-subsystem realization with *K*_+_ = *a*^†^*b*^†^, the pair amplitude satisfies ⟨*K*_+_⟩∝*N*_*A*_∝*M*_0_ when the subsystems contribute approximately equally. The approximately linear scaling observed in the experimental data therefore favors a bipartite interpretation over collective single-manifold squeezing.

## Acknowledgments

The author acknowledges the use of ChatGPT (OpenAI, GPT-5.4 Thinking), Claude (Anthropic), and Google’s Gemini as AI assistants for structural brainstorming, language refinement and translation, and LaTeX typesetting during the drafting of this manuscript. The author bears full responsibility for the accuracy and originality of the scientific arguments and equations presented herein.

